# Sustained exposure to high dietary ethanol impairs bumblebee survival and locomotor performance

**DOI:** 10.64898/2026.09.29.749860

**Authors:** Daniel Bajorek, Weronika Antoł, Krzysztof Miler

**Author notes:** Correspondence: Krzysztof Miler.

## Abstract

Ethanol occurs naturally in sugar-rich foods used by insects, but its effects in bees have been studied primarily in honeybees. We tested whether dietary ethanol affects survival and locomotor performance in workers of the bumblebee *Bombus terrestris* (L.). In the survival experiment, workers were fed ethanol (0, 1, or 10%) for 21 days; in the locomotor experiment, they were fed the same doses for one or seven days, after which we measured spontaneous movement using an open-field assay. We measured the proportion of time in movement, speed of movement, path meander (curviness), and arena exploration. The 10% ethanol concentration greatly increased the risk of mortality, whereas the estimated effect of 1% ethanol was smaller and statistically inconclusive. In terms of locomotion, the largest differences occurred after seven days of exposure to 10% ethanol. Workers in this treatment showed greater path meander and lower exploration than control workers. Speed was also lowest in this group. Evidence that exposure for seven days was worse than for one day at the same concentration was detected only for exploration. No clear locomotor effect was detected at 1% ethanol exposure. These findings provide comparative data on sustained ethanol exposure in bumblebees and show a strong concentration-dependent contrast, with substantial effects under high-concentration challenge but no clear effects on the measured endpoints at 1%.

## Introduction

The diets of animals are complex chemical mixtures. Plant-based food resources contain secondary metabolites as well as compounds produced by microorganisms that colonize them (Martin et al., 2022; Stevenson, 2020). Ethanol is one such microbial product (Lievens et al., 2015). Yeasts and other microorganisms produce ethanol when fermenting sugar-rich substrates, including fleshy fruits, plant sap and floral nectar (Bowland et al., 2025; Dudley and Maro, 2021). A recent survey detected ethanol in nectar from a wide range of plant taxa, although concentrations in positive samples were low, ranging from approximately 0.008 to 0.056%, with a mean close to 0.016% (Maro et al., 2026). These levels of ethanol may appear low, but they may still be biologically meaningful for small nectarivores foraging on large amounts of nectar relative to their body mass (Maro et al., 2026). Natural occurrence therefore provides a basis for studying how nectar-feeding insects respond to ethanol, while experimental exposure at substantially higher concentrations can address a different question: how these insects respond when dietary ethanol becomes a physiological and behavioral challenge. These contexts need to be distinguished because ethanol effects depend on concentration, exposure duration, and the biological endpoint considered (Bowland et al., 2025; Dudley and Maro, 2021; Miler, 2025).

Responses to ethanol can vary with concentration, exposure history, and what is analyzed. Mild stress sometimes produces a biphasic response, with improvement in some traits at a lower dose and inhibition at a higher dose (Agathokleous, 2018; Calabrese, 2014; Schirrmacher, 2021). Such responses are often trait-specific: a concentration eliciting little effect or apparently beneficial in one trait may nevertheless come at the cost of other traits, including survival, reproduction, immune function, or motor performance (Berry and López-Martínez, 2020; Cutler et al., 2022). Therefore, dietary ethanol exposure can be viewed as a stressor whose effects oscillate between benefit, tolerance, challenge, and impairment (Miler et al., 2018; Ostap-Chec et al., 2025b). Duration of exposure and its frequency are especially important in this context. Short-term access can reveal intoxication-like effects, whereas repeated exposure can produce cumulative damage, physiological adjustment, functional tolerance, or withdrawal-like responses after exposure ends. In honeybees, ethanol has been reported to alter walking, posture, grooming, righting responses, flight kinematics, learning, odor discrimination, foraging decisions, and social communication (Abramson et al., 2000; Ahmed et al., 2022; Bozic et al., 2006; Maze et al., 2006; Mustard et al., 2008; Sokolowski et al., 2012). Repeated exposure has been associated with altered survival, cellular stress, DNA methylation, and withdrawal (Hranitz et al., 2010; Ostap-Chec et al., 2024, 2021; Rasmussen et al., 2021; Yang et al., 2025). Therefore, to understand the consequences of a given ethanol concentration, it is crucial to specify how long and how often it is available.

Locomotor performance represents a valuable endpoint for detecting sublethal effects since it reflects sensory, neural, muscular, and energetic function. Furthermore, reliable movement is necessary for nectar-feeding pollinators when foraging (e.g., flying direction), handling resources or brood care, and maintaining the nest. Honeybee studies show that ethanol may alter walking, balance, coordination, and flight kinematics, with clearer effects at higher concentrations (Abramson et al., 2000; Ahmed et al., 2022; Maze et al., 2006). Given the biological consequences of those changes, such as reduced foraging efficiency, compromised return to the colony, and altered social interactions or resource transfer among nestmates (Sokolowski et al., 2012; Stauch et al., 2024; Stephenson et al., 2021; Wright et al., 2012), ethanol-induced locomotory alterations may be interpreted as potential non-lethal functional impairment.

Most research on bees and ethanol has focused on the honeybee (Abramson et al., 2025). The only previous controlled study on bumblebees examined how 1% ethanol and floral color affected the behavior of *Bombus impatiens* (Jones and Agrawal, 2022). Ethanol did not produce a uniform response but changed color preference and the use of social information. Survival and basic locomotor performance during sustained dietary exposure have not been directly examined in bumblebees. Testing these endpoints in *Bombus* provides a way to assess how far patterns established mainly in the honeybee extend to another major bee lineage. Bumblebees are particularly useful for such comparative work because they rely heavily on carbohydrate-rich floral resources, can be maintained and tested experimentally, and are increasingly used as models in apidology (Lihoreau et al., 2025).

In this study, we asked whether *B. terrestris* workers show altered probability of survival or changes in locomotor performance when provided with ethanol in their diets. Workers received 0, 1, or 10% ethanol (v/v). The 1% treatment provided a lower experimental concentration that connected the study with previous bee research, whereas the 10% treatment served as a high-concentration challenge. In one experiment, survival was monitored over three weeks with continuous ethanol availability. In a separate experiment, locomotor performance was tested after either one or seven days of ethanol access. We measured the proportion of time in movement, speed during active movement, meander as an index of path tortuosity, and relative exploration. We predicted that 10% ethanol would increase mortality and impair locomotor performance. We expected smaller or undetectable effects at 1%, and stronger locomotor effects after seven days than after one day, particularly at 10%.

## Materials and Methods

### Animals and experimental diets

Separate sets of commercially reared *Bombus terrestris* (L.) colonies were used in the survival and locomotion experiments (Biobest, Belgium). Each colony was connected to a foraging arena. Active, uninjured workers were collected manually with forceps from these arenas.

Experimental diets were prepared from a 50% (w/v) aqueous Apikand solution (łysoń, Poland). Absolute ethanol (99.8% v/v) was added to obtain final ethanol concentrations of 0, 1, or 10% (v/v). Solutions were thoroughly mixed before use and provided *ad libitum* in gravity feeders.

### Experiment 1: Survival

Workers from three colonies were assigned to one of three diets: 0, 1, or 10% ethanol. Each colony contributed one cage to every dietary treatment, yielding nine cages in total. Each cage contained ten workers, and the experiment began with 90 workers. Bees were maintained at 21 °C for 21 days. Fresh diet was supplied daily. Mortality was recorded when the feeders were replaced, and dead workers were removed.

### Experiment 2: Locomotion

Workers from two separate colonies were used in the locomotion experiment. Each colony contributed one cage to each of five treatments: control diet for seven days, control diet for six days followed by 1% ethanol for the final 24 h, 1% ethanol for seven days, control diet for six days followed by 10% ethanol for the final 24 h, and 10% ethanol for seven days. The design comprised ten cages, one cage for every colony-by-treatment combination. Approximately 20 workers were initially maintained in each cage. Bees were kept at 21 °C. Diets were refreshed daily, and dead workers were removed.

On the final day, ten workers were recorded from each cage, giving 100 locomotor observations. Recording order was randomized. Each bee was gently placed in the centre of a circular Petri dish arena (14.5 cm diameter, 1 cm height) under a transparent lid. Spontaneous activity was recorded immediately for 5 min using a top-mounted camera (Sony, Japan) at a resolution of 1920 × 1080 pixels and 50 fps. Trajectories were extracted in AnimalTA (Chiara and Kim, 2023). We quantified the mean speed during active movement, meander (angular change per unit distance traveled) as an index of path tortuosity, and relative exploration of the arena. We also quantified the proportion of recording time spent moving.

### Statistical analysis

Analyses were performed in R (R Core Team, 2025). Figures were generated with the *ggplot2* package (Wickham, 2016). Model adequacy was evaluated in the *DHARMa* package (Hartig, 2024).

Survival was analyzed using a mixed-effects Cox proportional hazards model using the *coxme* and *survival* packages (Therneau, 2026, 2024). Individual time to death, measured in days, was used as the response. Individuals alive at the end of the experiment were right-censored. Dietary treatment was included as a fixed effect. Colony was included as a fixed blocking factor. Cage identity was included as a random intercept to account for dependence among workers maintained in the same cage. The overall contribution of treatment was evaluated using a likelihood-ratio comparison between the complete model and a reduced model containing colony and the cage random effect but not treatment. The proportional-hazards assumption was evaluated using scaled Schoenfeld residuals from a Cox model stratified by colony, and no evidence of violation was detected (χ^2^ = 1.28, df = 2, p = 0.53).

A supplementary cage-level analysis used restricted mean survival time through day 21 (Royston and Parmar, 2013). Cage-specific mean survival was calculated and analyzed with treatment and colony as fixed factors. Planned comparisons of the two ethanol treatments with the control were adjusted using the Holm method.

For locomotion analyses, each model included the five-level treatment factor (control; 1% for 1 day; 1% for 7 days; 10% for 1 day; and 10% for 7 days) and colony as fixed effects, as well as cage identity as a random intercept. Mean speed while moving and meander while moving were positive continuous variables and were analyzed using Gamma generalized linear mixed models with log links. Relative exploration was bounded between zero and one and was analyzed using beta regression with a logit link. These models were constructed with the *glmmTMB* package (Brooks et al., 2017). The overall contribution of treatment to each endpoint was evaluated by comparing the complete model with a reduced model retaining colony and cage but excluding treatment. Estimated marginal means were calculated for all five treatments on the response scale using the *emmeans* package (Lenth, 2025). To perform contrasts, Holm adjustment was used. Simulation-based residual diagnostics showed no evidence of residual non-uniformity, overdispersion, or problematic outliers for speed, meander, and exploration models. The proportion of time spent moving was strongly concentrated near one; therefore, this endpoint was only summarized descriptively.

## Results

### Experiment 1: Survival

The analysis included 90 workers: 74 died during the experiment, and 16 were right-censored on day 21. All 30 workers receiving 10% ethanol died during the experiment, compared with 24 of 30 workers receiving 1% ethanol and 20 of 30 control workers.

Dietary treatment affected survival overall (χ^2^ = 24.21, df = 2, p < 0.001; Fig. 1). Workers receiving 10% ethanol had a substantially higher mortality hazard than control workers (HR = 18.45, 95% CI [8.46, 40.26], p < 0.001). The estimated hazard was also higher under 1% ethanol, although this effect was more uncertain and its confidence interval included no difference from the control (HR = 1.75, 95% CI [0.95, 3.23], p = 0.075).

**Figure 1.**
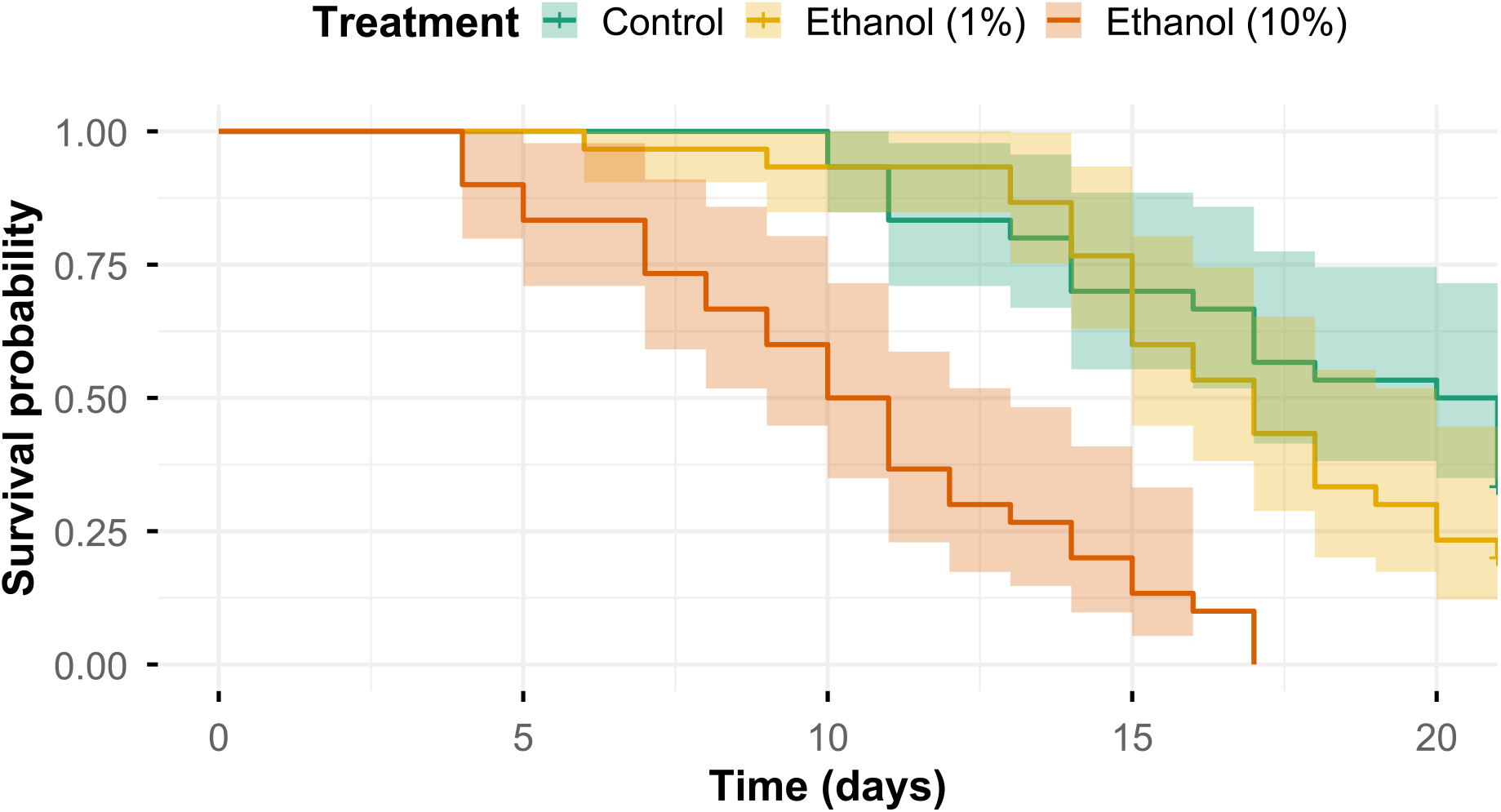
Survival of *Bombus terrestris* workers receiving control, 1% ethanol, or 10% ethanol diets. Kaplan-Meier curves show survival probabilities over 21 days. Shading represents 95% confidence intervals.

The supplementary cage-level analysis produced the same treatment pattern across all three colonies. Restricted mean survival through day 21 was 17.60 days in the control treatment, 16.70 days under 1% ethanol, and 10.43 days under 10% ethanol. Relative to the control, restricted mean survival was estimated to be 0.90 days lower under 1% ethanol (95% CI [−2.98, 1.18], p = 0.297) and 7.17 days lower under 10% ethanol (95% CI [−9.25, −5.08], p = 0.001). The reduction under 10% ethanol was present in each of the three colonies (Supplementary Table S1; Supplementary Figure S1).

### Experiment 2: Locomotion

Bees spent most of the recording time actively moving. Treatment means ranged from 0.931 to 0.967 (Supplementary Table S2; Supplementary Figure S2).

Treatment changed the speed of workers while moving (χ^2^ = 9.97, df = 4, p = 0.041; Fig. 2). The lowest estimated speed occurred after seven days of 10% ethanol (2.59 cm/s, 95% CI [2.26, 2.96]). The estimates for the other treatments ranged from 3.23 to 3.66 cm/s. However, speed after seven days of 10% ethanol was significantly lower only in comparison with speed after seven days of 1% ethanol (ratio = 0.71, 95% CI [0.59, 0.85], p < 0.001).

**Figure 2.**
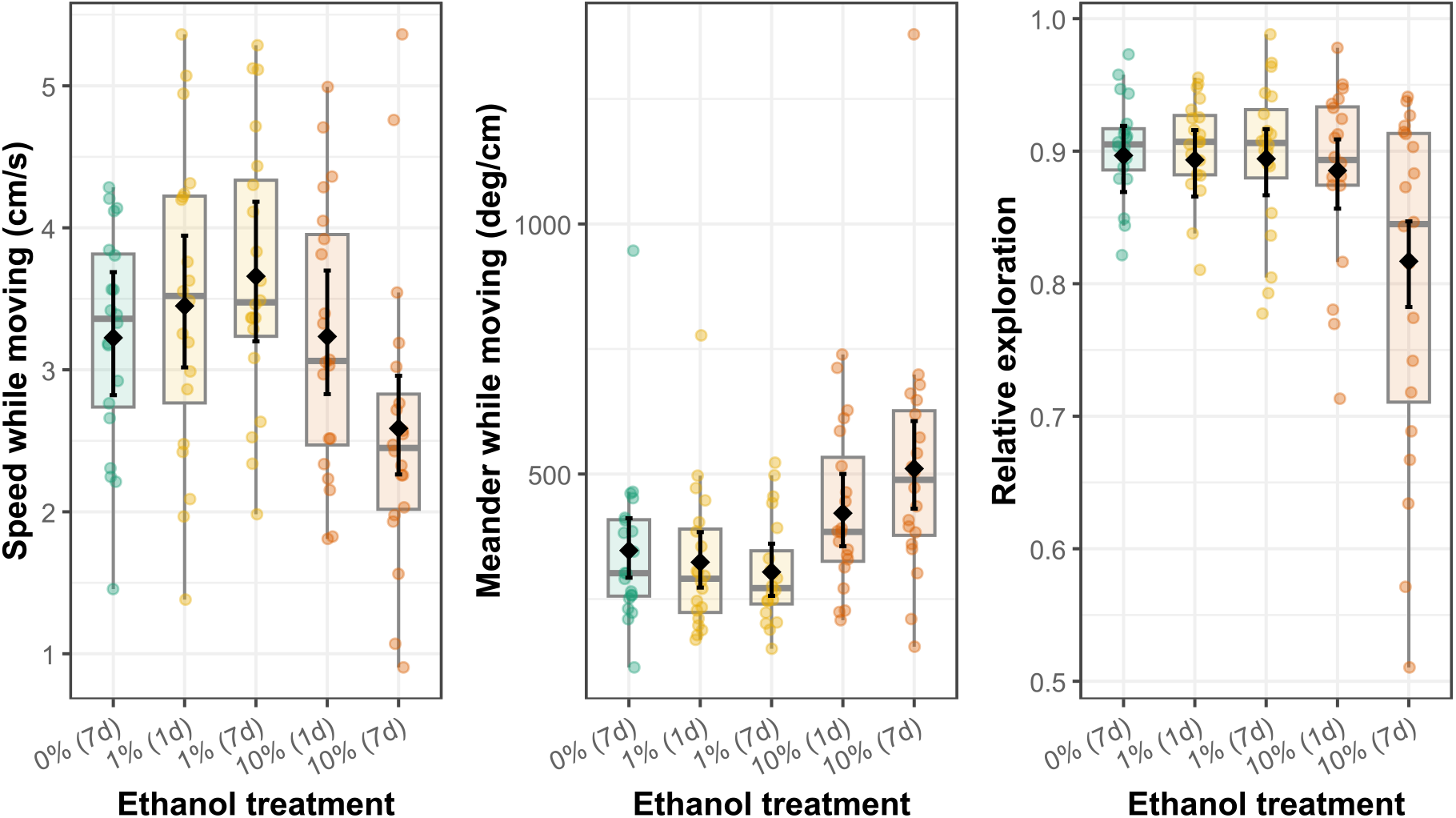
Locomotor performance of *Bombus terrestris* workers after exposure to dietary ethanol. Speed while moving (left panel), meander while moving (middle panel), and relative arena exploration (right panel) are shown for workers receiving the control diet or 1% or 10% ethanol for one or seven days. Colored dots show individual workers; boxes show interquartile ranges and whiskers extend to 1.5 × IQR. Black diamonds and error bars show model-estimated marginal means and 95% confidence intervals.

Treatment also affected meander while workers were moving (χ^2^ = 15.30, df = 4, p = 0.004; Fig. 2). The highest estimated meander occurred after seven days of 10% ethanol (511 deg/cm, 95% CI [431, 606]), compared with 347 deg/cm in the control treatment (95% CI [293, 412]). Thus, the workers showed approximately 47% greater meander than control workers (ratio = 1.47, 95% CI [1.16, 1.87], p = 0.010). In addition, meander was greater under 10% than under 1% ethanol after both one day (ratio = 1.30, 95% CI [1.02, 1.66], p = 0.032) and seven days of exposure (ratio = 1.68, 95% CI [1.32, 2.14], p < 0.001).

Relative arena exploration was affected by treatment as well (χ^2^ = 14.16, df = 4, p = 0.007; Fig. 2). Exploration was lowest after seven days of 10% ethanol, with an estimated value of 0.817 (95% CI [0.782, 0.847]). Estimates for the other treatments ranged from 0.885 to 0.897. Exploration after seven days of 10% ethanol was 0.080 lower than in the control treatment (95% CI [−0.120, −0.040], p < 0.001). It was also 0.068 lower than after one day of exposure to 10% ethanol (95% CI [−0.109, −0.027], p = 0.005). Thus, relative exploration was the only locomotor endpoint for which the direct comparison supported a stronger effect after seven days than after one day at 10%. Additionally, exploration after seven days of 10% ethanol was 0.077 lower than after seven days of 1% ethanol (95% CI [−0.118, −0.037], p < 0.001). No difference between the two concentrations was detected after one day of exposure.

Overall, the largest estimated locomotor changes occurred after seven days of 10% ethanol, but the strength of evidence differed among endpoints. This treatment was associated with greater path meander and lower exploration relative to the control. Speed was also lowest in this group, although it did not differ significantly from the control. Direct evidence that seven-day exposure produced a stronger response than one-day exposure at 10% ethanol was obtained only for relative exploration. No clear treatment-related differences were detected for the 1% ethanol groups.

## Discussion

Sustained dietary ethanol exposure produced a strong concentration-dependent contrast in *B. terrestris* workers. The high-concentration 10% challenge strongly increased mortality over 21 days, and locomotor differences were concentrated in workers exposed to this concentration for seven days. These individuals took more tortuous paths and explored less of the arena than control workers, and they also showed the lowest estimated speed. In contrast, the 1% treatment produced no clear difference in survival or in the analyzed locomotor measures relative to the control, although the survival estimate remained uncertain. The results therefore do not indicate generalized impairment across the tested ethanol treatments. Rather, they establish substantial effects under sustained high-concentration exposure alongside apparent resilience of the measured endpoints at the lower experimental concentration.

The survival result is consistent with the broader dose-related toxicity reported in honeybees. Bees offered a choice showed preferential consumption of 1.25-2.5% ethanol; however, concentrations of 2.5% or more increased mortality when ethanol-containing sucrose was the only diet (Mustard et al., 2019). Repeated 1% ethanol exposure shortened lifespan in relatively older individuals, but its effects were only observable after several days (Ostap-Chec et al., 2024; Rasmussen et al., 2021). In contrast, relatively younger workers showed no detectable survival cost during two weeks of forced 1% exposure or choice involving 0.5% ethanol (Ostap-Chec et al., 2025b). Infection might also shift this relationship. A survival advantage found with access to 0.625 or 1.25% ethanol during *Nosema* infection switched to a mortality increase and higher parasite loads at higher concentrations (Kuszewska, 2025). Parasitized bees increased ethanol preference too (Ostap-Chec et al., 2025a). Additionally, studies using ethanol as a solvent control show that low or moderate ethanol-only controls often had little effect on adult honeybee survival, whereas high ethanol concentrations were clearly harmful (Mura et al., 2020; Naree et al., 2025; Porrini et al., 2020; Tomé et al., 2023). Here, the analysis did not provide clear evidence of a survival effect at 1% ethanol, whereas the large estimated effect at 10% fits the broader pattern of high-dose toxicity.

The locomotor findings provide preliminary evidence that high-concentration ethanol exposure can alter selected aspects of bumblebee movement, in parallel with high-concentration effects in honeybees. Acute ingestion impairs walking, balance, grooming, and the righting response, while 1% can disrupt fine-scale motion (Abramson et al., 2000; Ahmed et al., 2022; Maze et al., 2006). Under repeated exposure, previous work demonstrated functional tolerance by showing reduced motor impairment after prior intoxication (Miler et al., 2018). It likewise increased resistance to sedation in young honeybees without inducing detectable alcohol dehydrogenase (ADH), indicating that functional tolerance cannot be attributed simply to increased ADH abundance (Miler et al., 2022). High-concentration exposure can disrupt gross locomotion in honeybees and was associated here primarily with greater path meander and lower exploration in bumblebees. Evidence that exposure for seven days produced a stronger effect than one-day exposure at 10% was detected for exploration, but not for speed or meander. It remains uncertain whether these changes emerge from neural toxicity, changed nutritional balance, reduced feeding, altered ethanol metabolism, or other processes. Our findings suggest that a distinction must be maintained between the physiological implications of acute intoxication and repeated ingestion of ethanol by pollinators.

The present study extends experimental research on ethanol beyond honeybees (Abramson et al., 2025). Other social Hymenoptera show substantial variation in ethanol effects. The Oriental hornet *Vespa orientalis*, for example, can tolerate extremely high amounts of ethanol, presumably through rapid metabolism and duplication of ADH genes (Bouchebti et al., 2024). Evidence from ants remains preliminary (Korczyńska et al., 2023). In bumblebees, Jones and Agrawal (2022) tested how 1% ethanol interacted with floral color to shape the behavior of *Bombus impatiens* workers. Ethanol did not produce a uniform response; instead, its effect depended on flower color and prior experience. The authors showed that ethanol can affect bumblebee foraging and social decisions, but only under specific sensory conditions (Jones and Agrawal, 2022). The present study addresses a different level of response. It found no clear change in the analyzed gross locomotor measures at 1%, even though subtler changes in decisions, sensory weighting, or social behavior may occur at that concentration. Indeed, an acute 1% exposure increased hemolymph ethanol without changing cognitive judgment bias or broad open-field locomotion in honeybees (Golański et al., 2026). The same concentration changed honeybee wing kinematics, including its pitch, heading, and stroke speed during tethered flight, but not gross performance measures (Ahmed et al., 2022). Two weeks of 1% access also left survival and flight endurance largely unchanged in relatively young honeybees (Ostap-Chec et al., 2025b). Together, these findings suggest that effects at lower experimental concentrations may become visible in specific decision-making contexts, social foraging situations, movement parameters, or fine-scale performance traits.

Several limitations define the scope of these conclusions. First, the 10% treatment was a high-concentration experimental challenge and should not be interpreted as representative of typical ethanol concentrations currently reported from floral nectar. The experiments therefore establish responses to sustained dietary ethanol across contrasting experimental concentrations rather than reproduce exposure under natural foraging conditions. Second, the experiments had limited independent replication, particularly at the colony level, so the absence of clear effects at 1% should not be interpreted as evidence that this concentration has no biological effects. We also cannot determine whether mortality and locomotor impairment at 10% arose mainly from ethanol toxicity, reduced feeding, altered energy balance, dehydration, metabolic costs, neural disruption, or a combination of these processes. The open-field assay provides controlled measures of spontaneous movement but does not directly measure flight, navigation, foraging success, or colony task performance. Future work using broader and more environmentally representative ethanol gradients, greater colony-level replication, and measurements of consumption, internal ethanol exposure, and physiological state will be needed to identify the mechanisms and ecological consequences underlying these responses.

In conclusion, dietary ethanol was associated with substantial mortality and changes in selected locomotor measures in *B. terrestris* under high-concentration challenge, whereas no clear survival or locomotor effect was detected at 1%. Repeated exposure to 10% ethanol was associated with higher mortality, greater path meander, and lower exploration; speed was also lowest after seven days at this concentration, although the evidence for this endpoint was less conclusive. These results extend experimental research on ethanol beyond primarily honeybee-based literature and provide a baseline for comparing ethanol responses among bee taxa. They also show that conclusions about ethanol effects in bees depend strongly on concentration, duration of exposure, and the biological endpoint examined.

## Acknowledgements

This work was supported by the National Science Centre, Poland [SONATA grant UMO-2021/43/D/NZ8/01044].

