## Supplementary Material for "Sustained exposure to high dietary ethanol impairs bumblebee survival and locomotor performance"

Correspondence:

**Table S1**. Cage-level survival outcomes and restricted mean survival time through day 21 in *Bombus terrestris* workers exposed to dietary ethanol. Each of the three source colonies contributed one cage of ten workers to each treatment. Restricted mean survival time through day 21 ($RMST_{(21)}$) was calculated as the cage mean of the observed survival or follow-up times, truncated at day 21.

| Colony | Cage | Treatment | Deaths | Right-censored | $RMST_{(21)}$ |
|---|---|---|---|---|---|
| **C1** | C1_C1 | Control | 8 | 2 | 16.4 |
| **C1** | C1_C2 | Ethanol 1% | 9 | 1 | 15.4 |
| **C1** | C1_C3 | Ethanol 10% | 10 | 0 | 7.4 |
| **C2** | C2_C4 | Control | 8 | 2 | 15.8 |
| **C2** | C2_C5 | Ethanol 1% | 10 | 0 | 15.0 |
| **C2** | C2_C6 | Ethanol 10% | 10 | 0 | 9.2 |
| **C3** | C3_C7 | Control | 4 | 6 | 20.6 |
| **C3** | C3_C8 | Ethanol 1% | 5 | 5 | 19.7 |
| **C3** | C3_C9 | Ethanol 10% | 10 | 0 | 14.7 |

Treatment affected $RMST_{(21)}$ overall ($F_{2,4} = 54.14$, $p = 0.001$). Model-adjusted $RMST_{(21)}$ was 17.60 days for the control, 16.70 days for 1% ethanol, and 10.43 days for 10% ethanol. Relative to the control, $RMST_{(21)}$ was estimated to be 0.90 days lower under 1% ethanol (95% CI [−2.98, 1.18], p = 0.297) and 7.17 days lower under 10% ethanol (95% CI [−9.25, −5.08], p = 0.001).

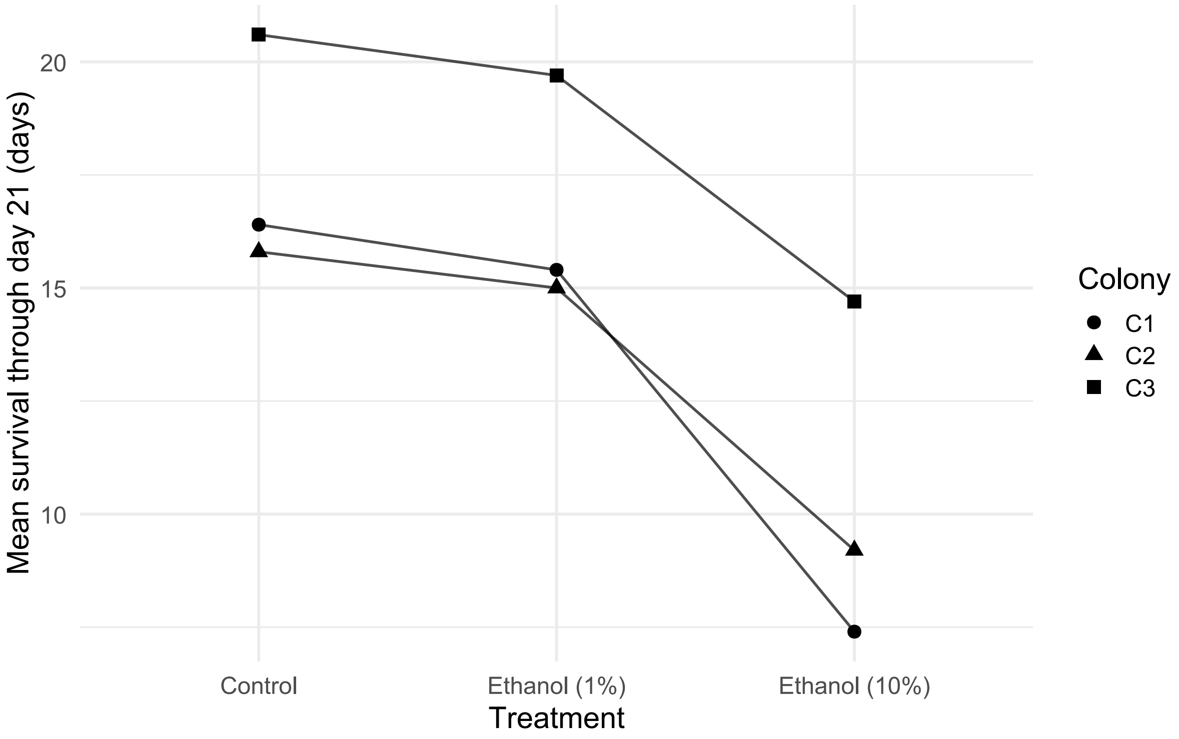


**Figure S1**. Cage-level restricted mean survival time through day 21 in *Bombus terrestris* workers exposed to dietary ethanol. Points show $RMST_{(21)}$ for each independently treated cage, and lines connect cages originating from the same source colony across the control, 1% ethanol, and 10% ethanol treatments. Symbol shape identifies the source colony. Each cage initially contained ten workers. $RMST_{(21)}$ was calculated as the mean observed survival or follow-up time truncated at day 21. The figure and analysis (Table S1 above) provide a cage-level robustness check for the pattern estimated by the individual-level mixed-effects Cox model.

**Table S2**. Descriptive summary of the proportion of recording time spent moving by *Bombus terrestris* workers exposed to dietary ethanol. Each treatment contained 20 recorded workers from two independently treated cages, with ten workers recorded from each cage.

| Treatment | Mean ± SD | Range |
|---|---|---|
| **Control** | 0.948 ± 0.057 | 0.784–0.998 |
| **1% ethanol, 1 day** | 0.967 ± 0.052 | 0.759–0.997 |
| **1% ethanol, 7 days** | 0.944 ± 0.081 | 0.740–0.996 |
| **10% ethanol, 1 day** | 0.958 ± 0.044 | 0.847–0.993 |
| **10% ethanol, 7 days** | 0.931 ± 0.098 | 0.648–0.997 |

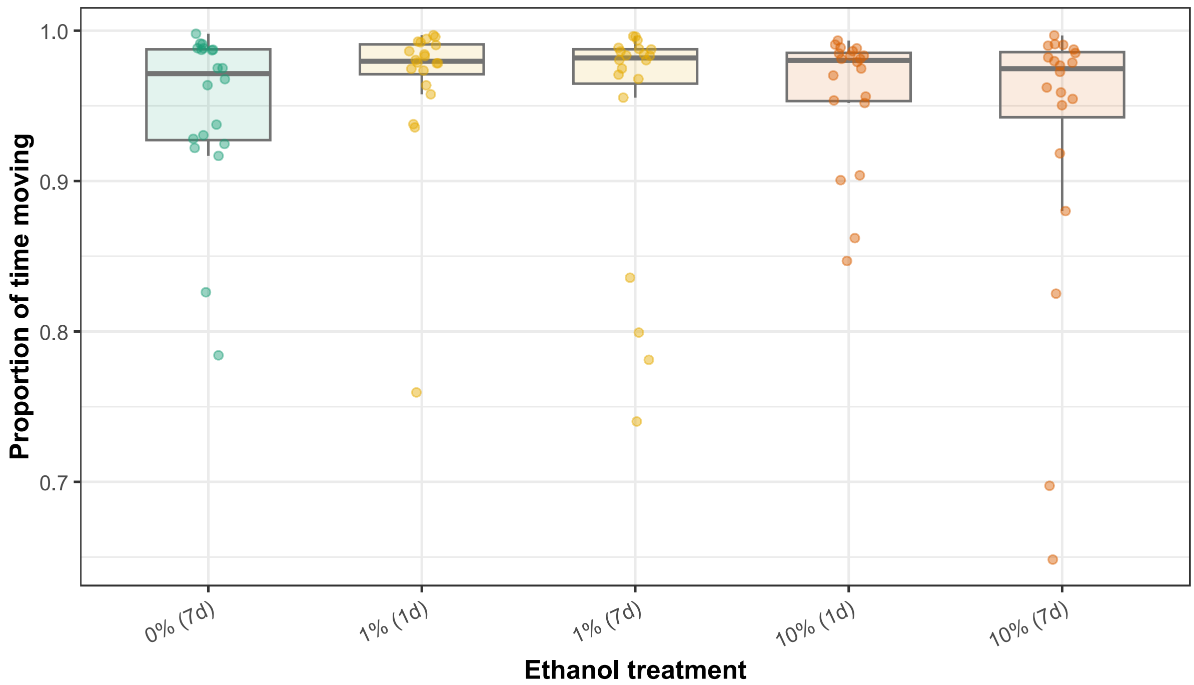


**Figure S2**. Proportion of recording time spent moving by *Bombus terrestris* workers after exposure to dietary ethanol. Colored points represent individual workers receiving the control diet or 1% or 10% ethanol for one or seven days. Boxes show interquartile ranges and whiskers extend to 1.5 × IQR.
